# Pancreatic Tumor-Derived Extracellular Vesicles Stimulate Schwann Cell Phenotype Indicative of Perineural Invasion via IL-8 Signaling

**DOI:** 10.1101/2023.06.26.546629

**Authors:** Emory Gregory, Isabel Powers, Azemat Jamshidi-Parsian, Robert Griffin, Younghye Song

## Abstract

Pancreatic cancer remains a pre-eminent cause of cancer-related deaths with late-stage diagnoses leading to an 11% five-year survival rate. Moreover, perineural invasion (PNI), in which cancer cells migrate into adjacent nerves, occurs in an overwhelming majority of patients, further enhancing tumor metastasis. PNI has only recently been recognized as a key contributor to cancer progression; thus, there are insufficient treatment options for the disease. Attention has been focused on glial Schwann cells (SC) for their mediation of pancreatic PNI. Under stress, SCs dedifferentiate from their mature state to facilitate the repair of peripheral nerves; however, this signaling can also re-direct cancer cells to accelerate PNI. Limited research has explored the mechanism that causes this shift in SC phenotype in cancer. Tumor-derived extracellular vesicles (TEV) have been implicated in other avenues of cancer development, such as pre-metastatic niche formation in secondary locations, yet how TEVs contribute to PNI has not been fully explored. In this study, we highlight TEVs as initiators of SC activation into a PNI-associated phenotype. Proteomic and pathway assessments of TEVs revealed an elevation in interleukin-8 (IL-8) signaling and nuclear factor kappa B (NFκB) over healthy cell-derived EVs. TEV-treated SCs exhibited higher levels of activation markers, which were successfully neutralized with IL-8 inhibition. Additionally, TEVs increased NFκB subunit p65 nuclear translocation, which may lead to increased secretion of cytokines and proteases indicative of SC activation and PNI. These findings present a novel mechanism that may be targeted for the treatment of pancreatic cancer PNI.

**Statement of Significance:** Identifying pancreatic tumor extracellular vesicles as key players in Schwann cell activation and perineural invasion by way of IL-8 will educate for more specialized and effective targets for an under-valued disease.

## 1. Introduction

The incidence of pancreatic ductal adenocarcinoma (PDAC) in the United States has continued to rise by 1% each year since 2000 and brings with it a bleak prognosis, notably an average 5-year survival rate of 11% across stages^1^. Although technology continues to advance, late-stage detection persistently increases patient mortality as symptoms only become discernable once metastasis has occurred^1^. In addition, within the last two decades, researchers have identified perineural invasion (PNI), in which cancer cells migrate into adjacent nerves to metastasize to secondary sites, as a key factor in the devastation of 70-95% of PDAC cases^2^. This attack on peripheral nerves contributes to elevated patient pain levels, while also increasing the likelihood of local tumor recurrence, even with complex surgical resection and adjuvant chemotherapy^2^. While personalized and targeted approaches to PDAC therapy have seen early clinical success, there are currently extremely limited treatment options for patients experiencing PDAC PNI^3^.

Cells within the tumor microenvironment (TME), including immune, stromal, and neural cells, regulate cancer cell activity including invasion. Evidence indicates that Schwann cells (SC) are one of the main drivers of PNI initiation and progression in PDAC^4^. These glial cells work to maintain the health of peripheral nerves, but upon injury or the onset of disease, they dedifferentiate from their mature state into an activated or repair phenotype. In the presence of cancer, SCs facilitate chemical gradients (e.g., tumor necrosis factor-alpha (TNF-*α*), interleukin 1 beta (IL-1*ββ*), IL-6, and CCL2) and produce cancer cell-mobilizing tracks (via matrix metalloprotease 2 (MMP-2), etc.) through the extracellular matrix (ECM) to advance PNI^5,6^. Additionally, SCs increase cell adhesion molecule presentation, e.g., L1 cell adhesion molecule (L1CAM), to bind cancer cells and physically mediate invasion^7^. Still, a clear mechanism by which SCs are activated in the TME remains unclear.

Alternatively, extracellular vesicles (EV) produced by tumor cells (TEV), including exosomes (diameter of 30-150 nm) and microvesicles (diameter of 100-1000 nm), have been implicated in cancer metastasis over longer distances than secreted factors alone^8^. These TEVs carry cargoes that promote ECM remodeling in secondary tissues and alter cellular behavior to form pre-metastatic niches primed for cancer colonization. In PDAC, TEVs instigate hepatic premetastatic niche formation by suppressing the immune response and have been shown to enhance cancer cell migration and proliferation in vitro^9,10^. Recent studies highlight cancer exosomes as contributors to tumor innervation, the relative opposite and common comorbidity of PNI^11^. While preliminary reports suggest pancreatic TEVs may activate SCs, the complete mechanism of this interaction and its role in PNI has yet to be explored^12^.

Among the known TEV-harbored cargoes, IL-8 is a potent factor that may promote SC activation and PNI. Circulating IL-8 has been linked to chemoresistance in PDAC, while pancreatic tumor microvesicle-derived IL-8 aids in the suppression of immune cells^13,14^. Recently, SCs have been found to heighten IL-8 production and nuclear factor kappa B signaling (NFκB) when activated by colorectal cancer cells; subsequently, cancer cells exposed to activated SC-conditioned media increased their migratory properties, evoking colorectal cancer PNI^15^. However, the contribution of PDAC TEV-derived IL-8 in PNI is not understood.

The role of the ECM in TEV function must not be overlooked. The ECM promotes biochemical and biomechanical signaling that allows for physiological cellular behavior^12^. To take advantage of this unique interaction, biomedical engineers have employed three-dimensional (3D) scaffolds created from natural or synthetic polymers to achieve mechanical and/or compositional similarity to the native tissues^11^. For example, Matrigel, a bioavailable ECM deposited by mouse sarcoma cells, has become a popular method for modeling the TME. Previous studies of PNI have implemented Matrigel to model the crosstalk between SCs and PDAC cells^4,5^; however, Matrigel and comparable products often vary batch-to-batch, have limited applications, and are unable to completely capture the physiological activity of cells within the TME^16^. More recently, tissue engineering has given researchers the ability to fabricate customizable biomaterials from native tissues through the process of decellularization^12^. This allows for more biomimetic replications of the TME. As such, we recently published a comparative analysis of peripheral nerve decellularization for SC-related disease modeling applications. Through this study, we were able to prove the value of a decellularized peripheral nerve ECM hydrogel in modeling quiescent and repair SC phenotypes^12^.

In this study, we implemented this platform to determine the role of pancreatic TEVs in reprogramming SCs in the peripheral nerve microenvironment to facilitate the progression of PNI. Here, IL-8 is found to be significantly elevated in TEVs over healthy cell EVs, regulates SC dedifferentiation into an activated and PNI-associated state, and potentially contributes to downstream NFκB signaling in SCs. We also highlight the efficacy of targeting TEV-derived IL-8 in limiting these phenotypic alterations in SCs. These data provide the first mechanistic proof of pancreatic TEV involvement in SC-induced PNI and a means by which to target PDAC PNI.

## 2. Materials and Methods

### 2.1 Cells

Human sNF96.2 Schwann-like, hTERT-HPNE (HPNE) immortalized pancreatic duct epithelial, and PANC-1 pancreatic ductal carcinoma cells were purchased from ATCC (Manassas, VA). HPNE cells were cultured in 75% glucose-free Dulbecco’s Modified Eagle Medium (DMEM; Thermo Scientific 11966025) and 25% M3 base (Incell M300) supplemented with 5% fetal bovine serum (FBS; Atlanta Biologicals S11150), 10 ng/ml human recombinant epidermal growth factor (Fisher Scientific PHG0311), 5 mM glucose (Fisher Scientific A2494001), and 750 ng/ml puromycin dihydrochloride (Fisher Scientific A1113803). PANC-1 cells were maintained in high glucose DMEM (Thermo Fisher 11965092) with 10% FBS and 1% penicillin-streptomycin (P/S; ATCC PCS999002). SCs were cultured in DMEM with 10% FBS, 1% P/S, 2 μM forskolin (Sigma F6886), 20 ng/ml neuregulin (Millipore 492028), and 1 g/L glucose.

### 2.2 EV Isolation

Upon reaching 80% confluency, HPNE and PANC-1 cells (passage number <20) were incubated with FBS-free DMEM for 24 hours. The next day, conditioned media were collected, and cells were passaged, noting the cell count. All passages contained ≥ 90% viability as quantified by trypan blue (Thermo Scientific 15-250-061). Conditioned media were centrifuged at 1000 rpm for 5 min to remove cell debris and then either added to Ultra-Clear Quick-Seal ultracentrifuge tubes (Beckman Coulter 246.344322), 100 kDa molecular weight cut-off centrifugal filters (MWCO; Sigma UFC910024), or reserved for ExoQuick Ultra kit isolation (Amsbio EQULTRA-20TC-1). Molecular weight filtered samples (MWF) were centrifuged at 2000 rpm until concentrated 10-fold. Kit-isolated EVs were collected according to ExoQuick Ultra (EQU) manufacturer’s instructions. Ultracentrifugation samples were balanced, added to the Beckman Coulter preparative ultracentrifuge 50Ti rotor, and processed at 200,000 g at 4°C for 90 min^17^. The resulting pellet was resuspended in phosphate buffered saline (PBS; VWR 97062-948) and ultracentrifuged at the same speed and temperature for 3 hours.

A portion of EVs was lysed according to previously established methods^18,19^. Briefly, for every 36 million cells of origin, 300 μl of 5% Triton X-100 (Sigma 93443) + 1% Halt protease/phosphatase inhibitor cocktail (Thermo Scientific 78440) were added to the harvested EVs. Samples were vortexed and sonicated at 25% pulse for 30 seconds. Samples were agitated on ice for 15 min, centrifuged at 14,000 g for 15 min, and quantified via BCA Protein Assay (Thermo Scientific 23210). All samples were stored at 4°C until further analyzed.

### 2.3 NanoSight Nanoparticle Tracking Analysis

EV samples were diluted appropriately and added to NanoSight NS300. Readings of each sample (n≥5) were conducted at 30-60 seconds per read using the continuous syringe flow. Results were adjusted according to the dilution factor.

### 2.4 Proteomic Analysis of EV Surface Markers and Cargoes

For proteomic analysis, PANC-1 and HPNE-conditioned media were ultracentrifuged 200,000 g at 4°C for 90 min, the pellets resuspended in PBS, and then re-centrifuged under the same conditions for 3 hours. Protein samples were reduced, alkylated, and digested using filter-aided sample preparation with sequencing grade-modified porcine trypsin (Promega)^20^. Tryptic peptides were then separated by reverse phase XSelect CSH C18 2.5 um resin (Waters) on an inline 150 mm x 0.075 mm column using an UltiMate 3000 RSLCnano system (Thermo). Peptides were eluted using a 60 min gradient from 98:2 to 65:35 buffer A:B ratio (i.e., buffer A containing 0.1% formic acid and 0.5% acetonitrile, buffer B containing 0.1% formic acid and 99.9% acetonitrile). Eluted peptides were ionized by electrospray (2.2 kV) followed by mass spectrometric analysis on an Orbitrap Exploris 480 mass spectrometer (Thermo). To assemble a chromatogram library, six gas-phase fractions were acquired on the Orbitrap Exploris with 4 m/z DIA spectra (4 m/z precursor isolation windows at 30,000 resolution, normalized AGC target 100%, maximum inject time 66 ms) using a staggered window pattern from narrow mass ranges using optimized window placements. Precursor spectra were acquired after each DIA duty cycle, spanning the m/z range of the gas-phase fraction (i.e. 496-602 m/z, 60,000 resolution, normalized AGC target 100%, maximum injection time 50 ms). For wide-window acquisitions, the Orbitrap Exploris was configured to acquire a precursor scan (385-1015 m/z, 60,000 resolution, normalized AGC target 100%, maximum injection time 50 ms) followed by 50x 12 m/z DIA spectra (12 m/z precursor isolation windows at 15,000 resolution, normalized AGC target 100%, maximum injection time 33 ms) using a staggered window pattern with optimized window placements. Precursor spectra were acquired after each DIA duty cycle.

### 2.5 Bioinformatics Analysis

Following proteomic data acquisition, data were searched using an empirically corrected library, and a quantitative analysis was performed to obtain a comprehensive proteomic profile. Proteins were identified and quantified using EncyclopeDIA and visualized with Scaffold DIA using 1% false discovery thresholds at both the protein and peptide level^21^. Protein exclusive intensity values were assessed for quality using ProteiNorm^22^. The data were normalized using VSN and statistical analysis was performed using linear models for microarray data (limma) with empirical Bayes (eBayes) smoothing to the standard errors^23,24^. Proteins with an FDR-adjusted p-value < 0.05 and a fold change > 2 were considered significant. EV surface protein markers were visualized using Scaffold DIA. Significant protein results were imported into QIAGEN Ingenuity Pathway Analysis (IPA), where relevant pathways were selected and their proteins visualized using Scaffold DIA. Proteins associated with IL-8 signaling were mapped via VolcanNoseR.

### 2.6 SDS-PAGE and Western Blot

EVs were lysed and then concentrated further with 3 kDa MWCO filters. Protein was quantified with BCA Protein Assays, and 10% (v/v) trichloroacetic acid (VWR BDH9310) was added to the lysed protein. Samples were centrifuged at 13,000 rpm for 3 min, rinsed with 100% acetone (VWR VWRVE646), and reconstituted in 8 M urea (VWR 97063-802). Laemmli buffer (Bio-Rad 1610747) with 10% (v/v) 2-mercaptoethanol (Bio-Rad 1610710) was combined 1:3 with the prepared protein and then incubated at 95°C for 3 min. Running buffer of 25 mM Tris base (Sigma 93362), 192 mM glycine (Sigma 410225), and 0.1% (w/v) sodium dodecyl sulfate (SDS; Sigma 75746) was added to the electrophoresis, and samples were loaded into 15% polyacrylamide gels. Electrophoresis was run for 60 min at 200 V. One gel was allocated for Coomassie blue staining (Bio-Rad 1610803), in which gels were stained for 45 min, rinsed in water, and de-stained overnight. Gels were imaged using the ChemiDoc Imaging System (Bio-Rad).

The transfer was conducted using the Trans-Turbo Transfer System with 0.2 μm nitrocellulose membranes (Bio-Rad 1704158). Antibody staining for albumin (1/1000; Cell Signaling 4929) was performed using the Pierce Fast Western Blot Kit according to the manufacturer’s instructions (Thermo Scientific 35050). Finally, the membrane was incubated in SuperSignal West Dura Extended Duration Substrate (Thermo Scientific 37071) for 4 min and then imaged on ChemiDoc using auto-optimal settings.

### 2.7 Decellularized Nerve ECM Hydrogel

Porcine sciatic nerves (Tissue Source LLC, Zionsville, IN) were decellularized as previously described^12^. Briefly, nerves frozen at -80°C were thawed at 37°C and rinsed in deionized water for 7 hours at room temperature (RT). Tissue was then subjected to the following washes at RT with agitation: 125 mM SB-10 (Sigma D4266) in 50 mM sodium/10 mM phosphate buffer (VWR BDH9286, BDH9298, and BDH9296; 18 hours); 100 mM sodium/50 mM phosphate buffer (VWR BDH9286, BDH9298, and BDH9296; 15 min); 3% (w/v) sodium deoxycholate (SD; Sigma D6750)/0.6 mM SB-16 (Sigma H6883) in 50 mM sodium/10 mM phosphate buffer (2 hours); 100 mM sodium/50 mM phosphate buffer (15 min x 3); SB-10 (7 hours); 100 mM sodium/50 mM phosphate buffer (15 min); SD/SB-16 (1.5 hours); and 50 mM sodium/10 mM phosphate buffer (15 min x 3). A 0.773% (w/v) magnesium chloride (VWR BDH9244) solution was created with 50 mM sodium/10 mM phosphate buffer, DNase (Sigma D4527) was diluted at 0.446% (w/v) in 0.15 mM sodium chloride, and a 75 U/ml DNase solution was made by combining the two resulting solutions. Nerves were incubated in the DNase solution for 3 hours at RT without agitation and then rinsed in 50 mM sodium/10 mM phosphate buffer with agitation (1 hour x 3, RT). ChABC (Sigma C3667) was diluted at 0.2 U/ml in PBS and used to wash the tissue for 16 hours at RT without agitation. Finally, nerves were rinsed thrice in PBS for 3 hours at RT, and the samples were stored dry at -20°C. Decellularized nerves were lyophilized for 3 days, chopped finely, and digested in a 1 mg/ml pepsin (Sigma P7000)-hydrochloride acid solution (Sigma 320331) at 12 mg tissue/ml pepsin-hydrochloride acid for 3 days at 400-500 rpm.

### 2.8 Platform Fabrication

Silicon wafers were patterned with cylindrical columns 200 μm tall and 4 mm in diameter (small) using photolithography as previously established^12^. Additionally, molds were 3D printed using acrylonitrile butadiene styrene with columns 200 μm tall and 8 mm in diameter (large). Polydimethylsiloxane (PDMS; DOW 1317318) microwells were fabricated using both molds, plasma-treated for 4 min, treated with 1% poly(ethyleneimine) (Sigma 181978; 10 min) and 0.1% glutaraldehyde (Sigma G6257; 30 min), and finally rinsed with sterile PBS before use.

### 2.9 EV Inhibition

Chilled nerve ECM pre-hydrogel was combined with 10% Medium 199 (Sigma M0650) and neutralized to pH 7.4 with 1 M sodium hydroxide (Sigma 415413). sNF96.2 cells (passage number <10) were cultured to 60-80% confluency, passaged, resuspended according to **Equation 1**, and added to the neutralized hydrogel at 1.5e6 cells/ml pre-hydrogel mixture. The mixture was added to small and large PDMS microwells, topped with flat PDMS lids, and incubated at 37°C for 30 min. Complete medium was added to the cultures, lids were removed, and the hydrogel cultures were maintained at 37°C for 1 day to regain physiological morphology. On day 2, freshly isolated EVs (50 μg/ml) ± anti-IL-8 neutralizing antibody (2 μg/ml; Novus Biologicals MAB208) were added to each 3D culture. Cultures were maintained for 2 days with media changes ± EV treatments/inhibition every 24 hours. Untreated SC-laden hydrogels acted as control cultures.

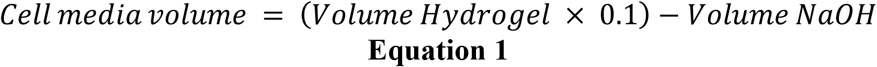

### 2.10 Cell Metabolism

AlamarBlue Cell Viability Reagent (Fisher Scientific DAL1100) was added to fresh complete media 1:9. Culture media were replaced with 1 ml alamarBlue solution/well and then incubated away from light at 37°C for 3 hours. Following incubation, conditioned media were added to a black 96-well plate, and the fluorescence was read on BioTek Synergy Mix Microplate Reader with an excitation/emission of 504 nm/531 nm. Percent difference in metabolism was calculated according to **Equation 2**, with R_C_ as the fluorescence of the alamarBlue reagent incubated without cells and R_S_ as the sample fluorescence.

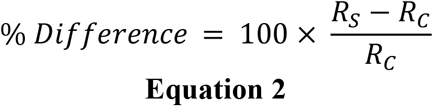

### 2.11 Immunofluorescence

Previously established protocols were implemented for the immunofluorescent staining of samples^12,25^. Briefly, cultures in small microwells were incubated in 3.7% formaldehyde at 4°C for 1 hour. For L1CAM (1/1000; Abcam ab24345) and glial fibrillary acidic protein (GFAP) staining (1/1000; Dako Z033429-2), fixed samples were rinsed at RT with PBS, permeabilized with 0.05% Triton X-100 in PBS (5 min), and blocked with 1% bovine serum albumin (BSA) in PBS with gentle agitation (1 hour). Domes of primary antibody dilution in 1% BSA were added atop each culture, and samples were incubated at 4°C (overnight). The following day, samples were rinsed at RT with 0.05% Tween-20 in PBS (5 min x 3) and incubated with secondary antibodies (1/500; Thermo A11029, A11029) and DAPI (1/2500; Fisher D1306) in 1% BSA away from light (1 hour). For NFκB p65 probing, fixed samples were permeabilized at RT with 0.1% Triton X-100 (10 min), blocked with 3% BSA (30 min), and incubated with primary antibodies (2.5 μg/ml; Abcam ab16502) in 3% BSA at 4°C (overnight). Hydrogels were rinsed at RT in PBS (5 min x 3) and then incubated with secondary antibodies in 3% BSA away from light (1 hour). All samples were stored in fresh PBS and imaged with the Olympus IX-83 inverted confocal microscope using 20X magnification and 2X digital zoom.

### 2.12 Luminex Multiplex Analysis

For EV cargo analysis, lysed EVs were concentrated with 3 kDa MWCO centrifugal filters and characterized via Luminex Multiplex Assay for TNF-*α* and IL-8 according to the manufacturer’s instructions (R&D Systems). Results were normalized by total protein content. On day 3 of the culture, SC-laden hydrogels were incubated in serum-free media overnight. The conditioned media was concentrated with 3 kDa MWCO filters and analyzed for TNF-*α*, IL-1β, IL-6, and IL-8 according to the manufacturer’s instructions. SC cultures were evaluated for DNA content (Qiagen 69506, Promega E2670), and results were used to normalize Luminex readings.

### 2.13 Image Analysis

GFAP and L1CAM immunofluorescence images were imported into Fiji ImageJ, and Z-stacks were compressed to max intensity via Z-Project. Images were auto-thresholded, and mean channel intensities were quantified via RGB Measure Plus and normalized by nuclei count. Debris was removed from the background of figure images using a custom MATLAB code. A custom MATLAB code isolated nuclear and cytoplasmic intensities of NFκB immunofluorescence images, which were normalized by area. The ratio of nuclear to cytoplasmic protein intensity was then computed.

### 2.14 Statistical Analysis

T-tests, ordinary analyses of variance (ANOVA), two-way ANOVAs, and multiple comparisons tests (e.g., Tukey’s) were performed using GraphPad Prism 9.4. Significant outliers were identified and removed using the GraphPad Prism Outlier Calculator.

## 3. Results

### 3.1 The isolation of EVs using MWF is confirmed

Samples were isolated via ultracentrifugation, MWF, or EQU and then assessed for particle size distribution using NanoSight Nanoparticle Tracking Analysis (**Figure 1A**). MWF and EQU methods collected much fewer particles than ultracentrifugation, yet MWF size distributions of both EVs and TEVs were more similar to those of ultracentrifuged samples compared to EQU-harvested EVs. As a more accessible alternative to ultracentrifugation, MWF will be used for all future isolations of EV samples.

**Figure 1.**
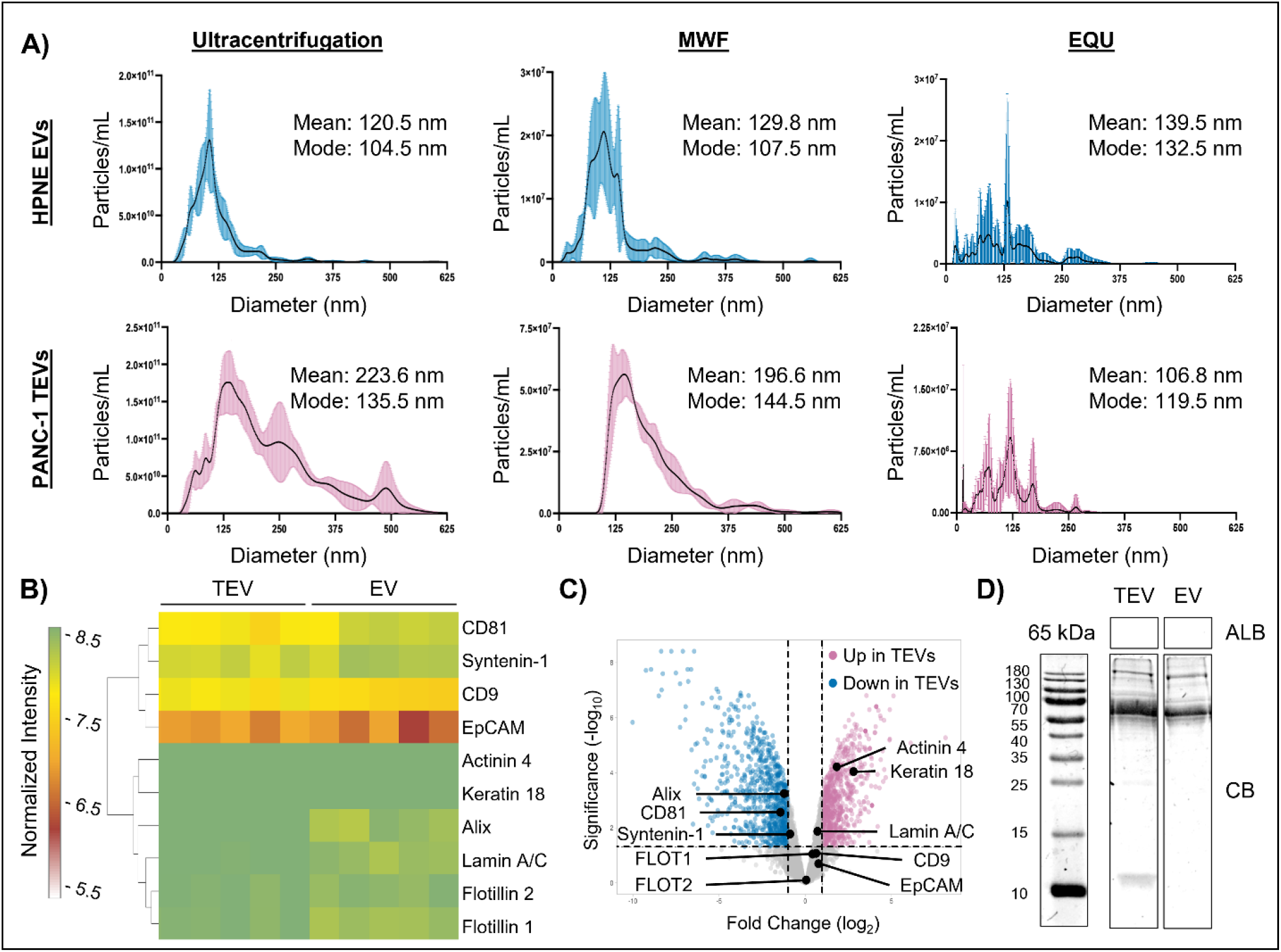
: EV isolation and characterization. A) Size distribution of ultracentrifuged EVs versus EVs isolated via molecular weight filtration (MWF) and ExoQuick Ultra (EQU). Validation of HPNE EV and PANC-1 TEV isolation (B-C) and absence of non-EV proteins (D).

Proteomic analysis of EVs and TEVs was then conducted to determine their potential role in PNI. These results were also used to confirm the presentation or absence of EV markers, which must include a transmembrane protein (group 1), a cytosolic marker with the ability to bind lipids or membranes (group 2), and a non-EV marker to prove a lack of cell culture contamination (group 3)^26^. Transmembrane proteins CD9 and epithelial cell adhesion molecule (EpCAM) were not statistically different in the samples, while CD81 was significantly elevated in EVs versus TEVs (**Figure 1B-C**). Of the group 2-classified proteins, flotillins 1 and 2 as well as syntenin-1 were found at similar levels in both types of EVs, but ALG-2-interacting protein X (Alix) was again elevated in HPNE samples. Additionally, cytoskeletal actinin 4 and keratin 18 were increased in TEVs, indicating the presence of large EVs^26^. Although one other large EV marker lamin A/C was not statistically different in the samples, these results verify the NanoSight analyses that showed larger EVs in PANC-1-isolated samples overall. Finally, a Western blot for albumin was performed and produced negative results (**Figure 1D**), confirming that particles seen with NanoSight were not contaminated by cell culture reagents.

### 3.2 IL-8 signaling-associated proteins are significantly elevated in pancreatic TEVs

Proteomics analysis was revisited to examine cargoes that may produce an activation response in SCs. The results highlighted 1550 differentially expressed proteins, with 815 proteins decreased and 735 proteins increased in PANC-1 TEVs versus HPNE EVs. These differentially expressed proteins then underwent pathway analysis to further identify potential targets for PNI (**Figure 2A**). When doing so, IL-8 signaling was ranked among the highest significance scores (p<0.0001) with a z-score of 0.73, indicating an elevation in TEVs. Proteins involved in IL-8 signaling were then visualized via heatmap (**Figure 2B**) and volcano plot (**Figure 2C**). While most of these elevated proteins were upstream regulators of IL-8 synthesis, PI3K-beta and NFκB subunits p50 and p65, which are both downstream products of IL-8 intake, were shown to be significantly higher in TEVs (p<0.05). Interestingly, NFκB inhibitor IkappaBbeta (IκBβ) was also diminished in TEV samples (p<0.05). IL-8 content finally was probed to determine if targeting the cytokine would be effective (**Figure 2D**). HPNE EVs were found to contain 7.09 ± 3.99 pg IL-8/ng of total protein, while TEVs showed levels of 74.07 ± 54.65 pg/ng protein (p<0.01). While IL-8 concentrations in both TEVs and EVs varied between cell passages, even the lowest TEV levels were twice that of the highest EV concentrations.

**Figure 2.**
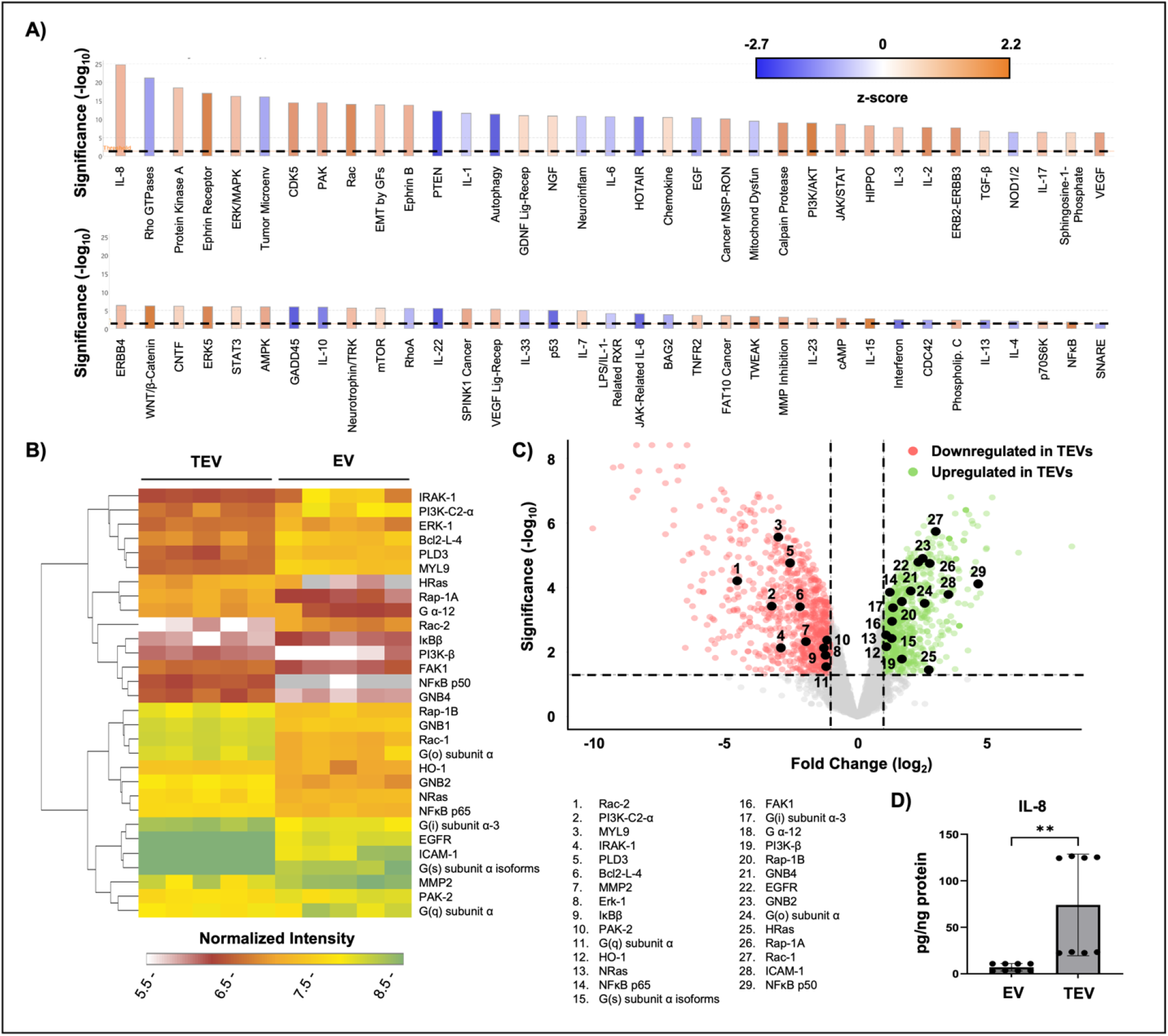
: EV protein and pathway analysis. Pathway analysis was conducted using Qiagen IPA (A). Proteins related to IL-8 signaling are indicated in the heatmap (B) and volcano plot (C). Validation of IL-8 content in EVs and TEVs (D). **p<0.01; significance (-log_10_) ≥ 1.3, and fold change (log_2_) ≥ |1| are significant.

### 3.3 Pancreatic TEVs activate SCs via IL-8

While compelling, to effectively determine the role of IL-8 signaling in SC activation and PNI, TEV- and EV-associated IL-8 was neutralized, and SCs were cultured in a biomimetic microenvironment with the resulting samples. Total IL-8 secreted by SCs was analyzed (**Figure 3A**), which highlighted a significant increase in IL-8 production by the EV (77.33 ± 1.03 pg/ng DNA) and TEV treatment groups (167.62 ± 4.07 pg/ng DNA) compared to untreated control samples (58.05 ± 1.56 pg/ng DNA; p<0.0001). Moreover, TEV-treated samples secreted significantly higher levels than EV-treated SCs (p<0.0001). Following IL-8 inhibition, the control (3.89 ± 0.02 pg/ng DNA), EV-treated (7.21 ± 0.20 pg/ng DNA), and TEV-treated SCs (1.17 ± 0.01 pg/ng DNA) experienced a reduction in IL-8 secretion (p<0.0001). These findings confirmed the aforementioned proteomics results as well as the uptake of EVs by SCs.

**Figure 3.**
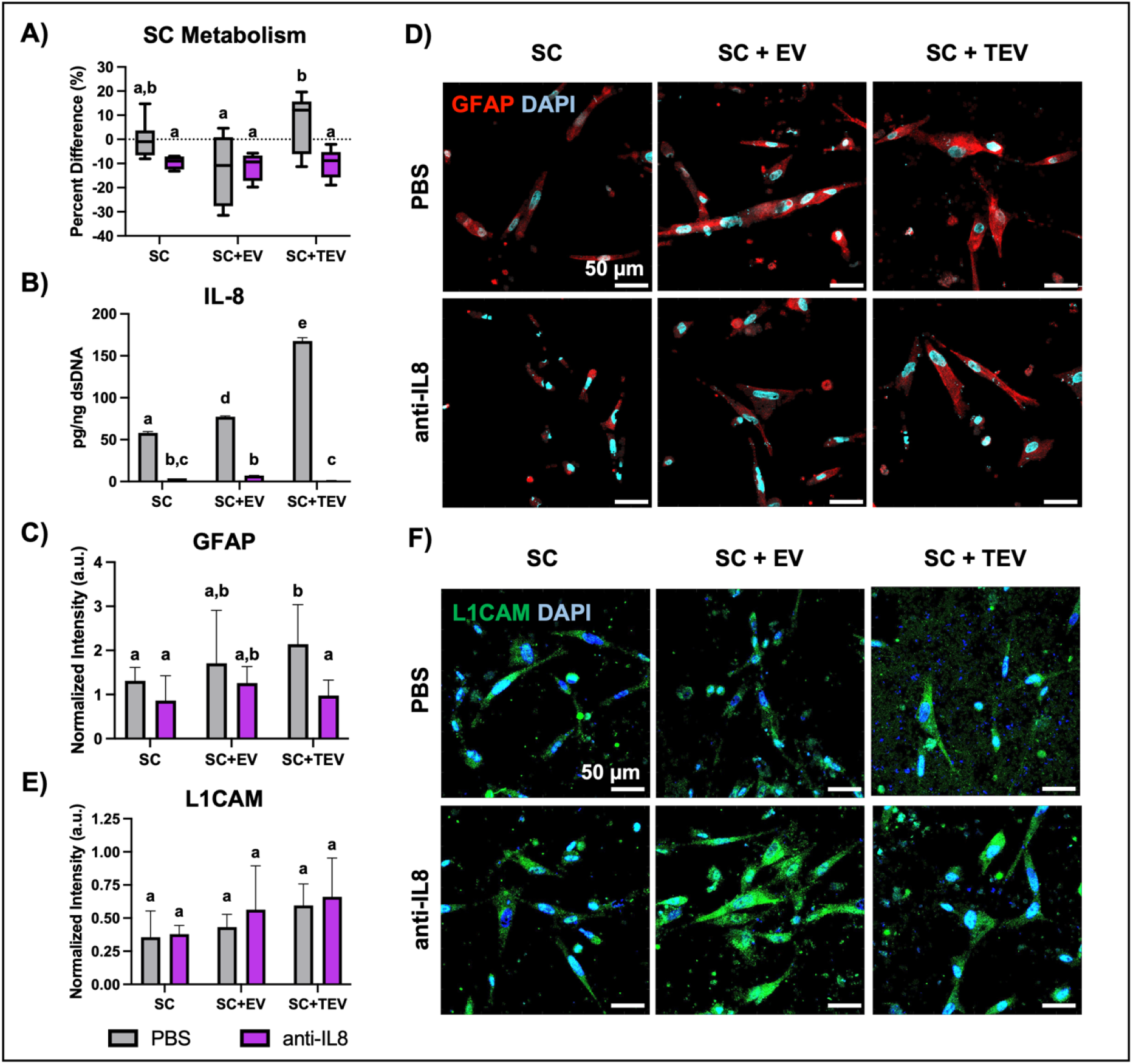
: Phenotypic changes via IL-8. IL-8 is neutralized following antibody inhibition (A), which alters SC metabolism (B) and GFAP levels (C-D). L1CAM levels remain unaffected (E-F). Shared letter designations indicate no statistical difference.

In addition, the effect of EVs and TEVs on SC metabolism was examined (**Figure 3B**). TEV-treated samples exhibited significantly higher metabolic activity that SCs treated with healthy EVs (p<0.005), which was stifled by IL-8 inhibition (p<0.01). Likewise, though not significant, SC metabolism dipped in control and EV-supplemented samples when IL-8 was neutralized. Treated groups were then evaluated for relative GFAP levels to confirm the dedifferentiation of SCs by TEVs (**Figure 3C-D**). TEV-supplemented SCs showed significantly higher GFAP intensities than those of untreated samples (p<0.05). This increase in activation levels was then depleted following IL-8 inhibition (p<0.01). However, while GFAP intensity also fell when IL-8 was inhibited in EV-treated and control cells, these changes were not as substantial nor significantly different. Although an increase in GFAP is indicative of activation, SCs treated with TEVs did not experience a shift in L1CAM nor were the levels depleted with IL-8 inhibition (**Figure 3E-F**). This suggests that TEVs and IL-8 may not contribute to a change in cell adhesion molecule presentation and promote PNI in this manner.

### 3.4 Pancreatic TEVs may induce NFκB signaling to prompt SC phenotypic shift

Based on the pathway analysis, NFκB p65 was probed after determining that pancreatic TEVs do promote SC dedifferentiation via IL-8. Immunofluorescence showed a significant increase in the sequestering of p65 to the nucleus in SCs treated with TEVs compared to control SCs (**Figure 4A-B**). SC secretome was evaluated for that associated with NFκB signaling (**Figure 4C-G**), which found that TEV-supplemented SCs secreted statistically higher levels of IL-1β, TNF-*α*, IL-6, and CCL2 over control (p<0.0001) and EV-treated SCs (p<0.0001). While the elevated concentrations of IL-1β, TNF-*α*, and IL-6 were neutralized in all treatment groups following IL-8 inhibition, CCL2 only decreased in TEV-supplemented cells. MMP-2 was also secreted at higher concentrations than both control (p<0.0001) and EV-treated groups (p<0.005). However, IL-8 inhibition only significantly affected MMP-2 in TEV-treated SCs (p<0.0001), while control and EV-treated groups increased their secretion levels (p<0.0001). Proteomics found that MMP-2 was significantly elevated in HPNE EVs (not shown), which may contribute to this relationship. Given these findings, IL-8/NFκB signaling will continue to be investigated for their contributions to SC activation.

**Figure 4.**
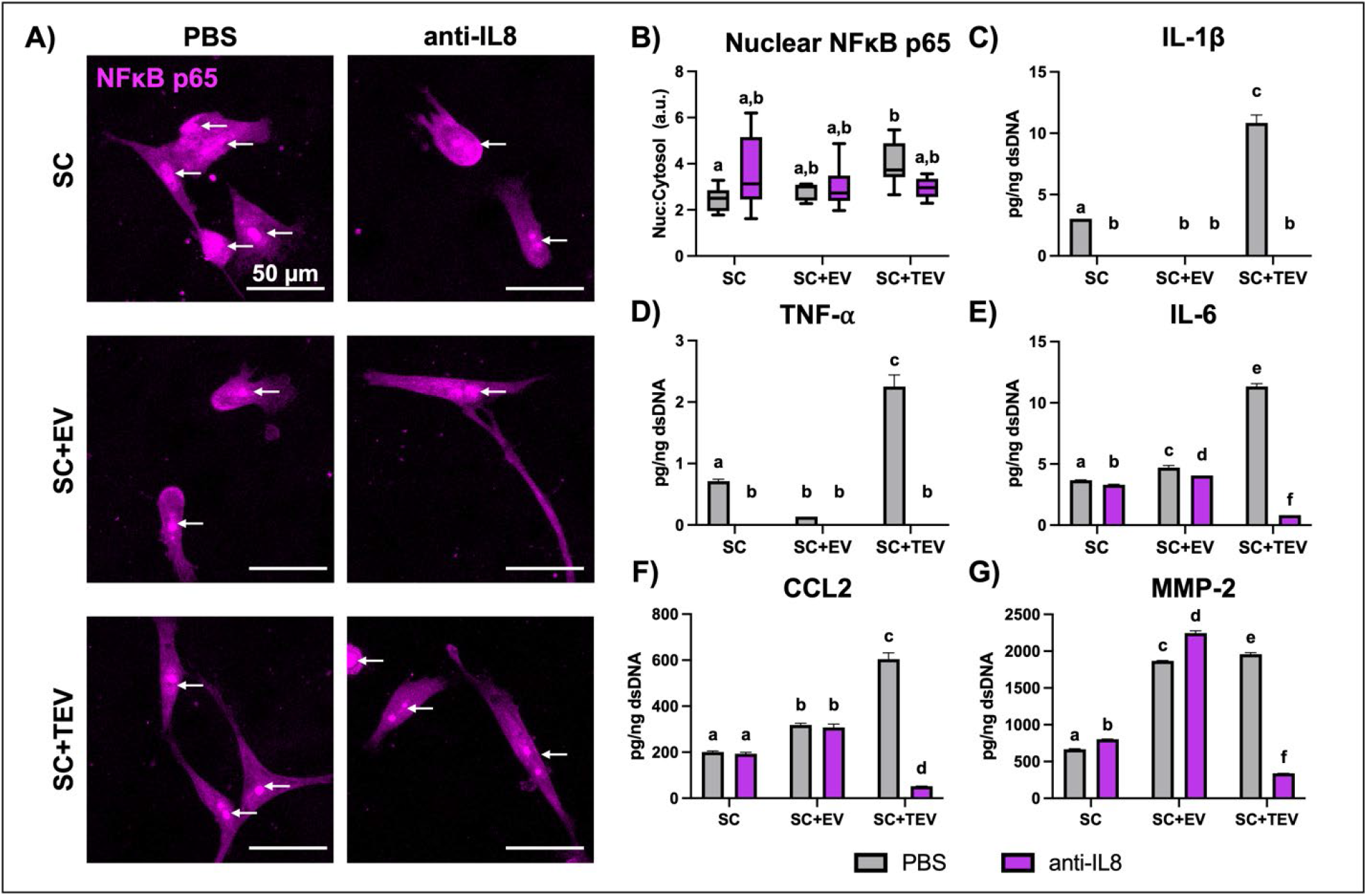
: NFκB signaling. SCs treated with TEVs sequester NFκB p65 in the nucleus (A-B). Such signaling induces the secretion of IL-1ββ (C), TNF-α (D), IL-6 (E), CCL2 (F), and MMP-2 (G). Shared letter designations indicate no statistical difference.

## 4. Discussion

In the present study, we explored the relationship between TEVs and SCs in PDAC PNI. We found that TEVs contain increased levels of IL-8 and proteins related to IL-8 signaling, including NFκB p65. TEVs were able to dedifferentiate SCs into an activated state via IL-8, resulting in the nuclear translocation of NFκB p65 and the production of IL-8, IL-1β, TNF-*α*, IL-6, CCL2, and MMP-2. However, the inhibition of IL-8 in healthy cell-derived EVs did not necessarily lead to the same effect. This suggests that IL-8 plays a unique role in cancer and may be a viable option for PNI-targeted therapies.

In the short time frame in which TEVs have been studied, researchers have put a major focus on understanding exosomes in particular. More recently, however, larger EVs have been embraced as significant players in cancer progression. In comparing EV markers, PANC-1 TEVs were found to have higher concentrations of actinin 4 and keratin 18 than HPNE EVs, which the International Society for Extracellular Vesicles recognize as large EV markers (e.g., microvesicles)^26^. In addition, Sun et al. found that microvesicles harbor higher levels of Alix and CD81 compared to exosomes; PANC-1 TEVs isolated in this study present in the same manner^27^. These results in combination with NanoSight readings suggest that PANC-1s produce higher levels of microvesicles while HPNE EVs are mostly exosomes. Santana et al. compared healthy and cancer epithelial cell-derived EVs from breast, brain, and pancreatic cell lines and presented similar findings^28^. Not only was this trend seen in PANC-1 TEVs but also those isolated from lower-stage BxPC3 cells^28^. Further examination of EVs from pancreatic cancer cells at various stages may help determine clinical translatability of our findings.

Pathway analysis of proteomics results found that IL-8 signaling is among the highest signaling pathways upregulated by TEVs over EVs. IL-8 is a broad prognostic factor in many cancers, including PDAC^29^. Additionally, IL-8 promotes a stem-like phenotype, proliferation, and chemoresistance in cancer cells as well as angiogenesis^29,30^. Because IL-8 also recruits immune cells under pathological conditions, recent studies have explored neutralizing IL-8 through targeted approaches and combinational therapy with immune checkpoint inhibitors^31,32^. While it is clear that activated SCs secrete IL-8 under PNI, there is limited evidence of the effects of IL-8 on SCs^33^. Chen and Chen recently highlighted that colorectal cancer cells induce IL-8 production in SCs via NFκB and that inhibiting IL-8 decreased SC activation, which we also found to be in agreement^15^. The present study is unique as it validates IL-8 as an activator of SCs, which includes promoting the production of IL-8, IL-1β, TNF-*α*, IL-6, CCL2, and MMP-2. Even so, TEVs and IL-8 did not appear to affect L1CAM presentation by SCs, suggesting that TEVs may not be responsible for the cell adhesion properties of SCs. Further explorations into the role of TEVs in cell adhesion molecule upregulation during PNI, including neural cell adhesion molecule, should be conducted^4^. Nonetheless, this altered phenotype mirrors that of SCs during PNI^34^.

Furthermore, while NFκB induces the secretion of cytokines and proteases including IL-8, IL-1β, TNF-*α*, IL-6, CCL2, and MMP-2 in cancer and other inflammatory diseases, the effects of NFκB in SCs have been neglected over the last 15 years. It is known that NFκB induces cytoskeletal modification in SCs in which GFAP is upregulated^35^. In this study, NFκB p65 was sequestered in the nucleus at higher concentrations than untreated SCs, which resulted in significant secretion of IL-8, IL-1β, TNF-*α*, IL-6, CCL2, and MMP-2. Although targeting IL-8 diminished the production of these proteins, nuclear p65 only moderately fell with IL-8 neutralization. The most common NFκB dimer in SCs is the p65-p50 complex, yet limited research exists outlining the role of each subunit in SC function^34,36^.

Finally, TEVs were found to increase SC redox metabolism, which could be attenuated with IL-8 inhibition. In lung epithelial cells, IL-8 has been shown to alter metabolism through glutathione reduction^37^. High glucose levels in cases such as type-II diabetes also activate SCs, but this mechanism leads to SC apoptosis and is inhibited by downstream NFκB signaling^38^. This warrants further investigation into SC metabolism and PDAC PNI.

## 5. Acknowledgements

This work was supported by the NIH through the award number P20GM139768, PhRMA Foundation, Arkansas Biosciences Institute, the University of Arkansas Women’s Giving Circle awarded to YHS, the University of Arkansas Honors College Research Grant awarded to IP, and IDeA National Resource for Quantitative Proteomics NIH/NIGMS grant R24GM137786. We would like to thank Drs. Sam Mackintosh and Stephanie Byrum for their work in conducting proteomic analysis and bioinformatics, respectively. We thank Drs. Jorge Almodóvar, Kartik Balachandran, Yuchun Du, Christopher Nelson, and Raj Rao for access to their equipment. We would also like to thank Amanda Walls for aiding in the fabrication of our 3D culture platforms.

